# Recording and reproducing the diurnal oviposition rhythms of wild populations of the soft- and stone- fruit pest *Drosophila suzukii*

**DOI:** 10.1101/342535

**Authors:** Bethan Shaw, Michelle T. Fountain, Herman Wijnen

## Abstract

*Drosophila suzukii* is a horticultural pest on a global scale which causes both yield and economic losses on a range of soft- and stone-fruit. Tackling this pest is problematic but exploiting behavioral rhythms could increase the impact of control. To do this, a better understanding of behavioral patterns is needed. Within this study we aimed to investigate rhythms in reproductive behavior of wild *D. suzukii* under natural conditions in the field. Environmental parameters were also recorded to decipher how they influence these rhythms. Assays were then performed on laboratory cultures, housed under artificial conditions mimicking the temperature and light cycles, to see if these patterns were reproducible and rhythmic. We were able to promote field like oviposition patterns within the laboratory using realistic temperature and light cycles regardless of variations in other factors including substrate, humidity, and lighting type. Locomotion activity was also recorded under these mimicked conditions to identify how this behavior interacts with oviposition rhythms. Both our field and laboratory assays show that oviposition behavior is likely under the control of the circadian clock and primarily occurs during the day, but these patterns will be disrupted by unfavorable temperature conditions. This was also found within locomotion rhythms. With an increased understanding of how these behaviors are influenced by environmental conditions, we highlight the importance of using realistic temperature and light cycles when investigating behavioral patterns. From an increased understanding of *D. suzukii* behavior we increase our ability to target the pest in the field.

## Introduction

The circadian clock is the internal daily pacemaker found within many different organisms that coordinates daily behavioural and physiological rhythms ^1^. Organisms from bacteria to mammals display a roughly 24-hour cycle which is synchronised to the rotation of the earth on its axis ^2,3^. Environmental time cues such as light and temperature entrain the clock and result in behavioural patterns occurring in synchrony with the environment and at optimum benefit to the individual ^4^. This results in individuals of the same species being synchronised to each other, allowing key behaviours such as mating or hunting to occur at the same time ^5,6^. This synchronicity provides an opportunity to exploit these behaviours.

Understanding behavioural rhythms for pest species is beneficial as control methods can be used to target behaviours, increasing the impact of control ^7^. One species whose behavioural patterns are of interest is the soft-, stone- and ornamental-fruit pest *Drosophila suzukii* (Matsumura). This pest causes global yield and economic losses as female *D. suzukii* cut into the epicarp of ripening fruit with a serrated ovipositor to deposit eggs. Once hatched, the developing larvae consume the fruit from within, resulting in collapse, rendering it unmarketable ^8–10^. With control measures having a limited impact and, those that do, disrupting integrated pest management practices, attention has turned to exploiting behavioural patterns for more effective control ^11^. One complication is that field behaviour of wild populations varies with regional and seasonal differences in factors such as light and temperature. Laboratory-based assays of rhythmic behaviours could be instrumental in predicting field behaviour as they are accessible year-round and offer superior experimental control. However, in both *D. suzukii* and *D.melanogaster*, a closely related and extensively studied model organism, field behaviour deviates from that observed in laboratory-based simulations ^11-16^. As egg-laying females are the source of the problem regarding *D. suzukii*, it is unsurprising that researchers have begun investigating patterns in oviposition ^17,18^. So far, these experiments have not focused on oviposition of wild populations in natural conditions or the ability to promote wild behaviour in a laboratory-based setting.

The aims of this paper were to (1) describe daily rhythms in wild *D. suzukii* reproductive oviposition in the field; (2) determine environmental parameters that regulate diurnal oviposition in *D. suzukii* and configure them to reliably recreate oviposition rhythms in the laboratory; and (3) determine the association between daily rhythms in locomotor activity and oviposition. Our results provide valuable insight into field-based reproductive oviposition rhythms of wild *D. suzukii* populations, which are relevant to integrated pest management strategies addressing this invasive species in horticulture. Moreover, by deducing how field oviposition behaviour might be reconstituted in the laboratory, we offer an approach that may be reversed to predict *D. suzukii* oviposition on fruit crops using a laboratory-based assay. Finally, we also probe the relationship of the oviposition and locomotor activity rhythms to reveal how these two behaviours are associated in the context of different environmental cycles.

## Materials and methods

### Field oviposition

Field oviposition rhythms were observed at times in the year to coincide with specific cherry crop events in the UK: cherry ripening in June (start date 21^st^ June 2016), cherry harvest in August (start date 02^nd^ August 2016) and end of cropping season in October (start date 02^nd^ October 2016). Oviposition assays were performed in an unsprayed, protected, strategic cherry orchard (cv. Sweetheart and Penny) located at NIAB EMR, Kent, UK. *D. suzukii* presence in the orchard was confirmed from DROSO TRAP^®^ monitoring traps from Biobest, baited with the Char- Landolt four component lure ^19^. These were deployed in the vicinity of the oviposition assay sites prior to commencing the trials. Captured females were dissected to determine fecundity at the beginning of each experiment by identification of mature ovaries and eggs ^20^. Traps were not removed from the orchard, but oviposition stations were deployed >30 meters away to reduce interference. Oviposition stations were created by modifying green delta traps (Agralan Ltd. UK). The open sides were covered with a 3 mm wire mesh to prevent entry of larger animals that may utilise the fruit. An entry door was added to allow access into the station via one of the solid sides. Stations were hung on cherry trees 1.5 m above the ground in shaded areas and spaced 15 m apart. Unsprayed ripe cherries, (kindly donated by G. H. Dean and Norton Folgate cv. Penny), or raspberry (kindly donated by Driscoll’s cv. Unknown) were washed, dried and then frozen for a minimum of 72 hours before use. Fruit was removed from the freezer two hours before deployment in the field to standardise fruit composition.

Due to insufficient field populations no reproductive oviposition occurred during the June field assay as so is excluded from the following field assay methods, results and analysis. See discussion for further details.

On commencing each experiment, either a single ripe cherry (August) or raspberry (October) was placed in the station within a Petri dish. Fruit type used was dependent on seasonal availability. After two hours, the fruit was removed from the station and transferred to a ventilated 70 ml specimen container (7 cm high × 5 cm diameter, polypropylene Sarstedt). A new piece of fruit was replaced in the station. Fruits were replaced within the station every two hours throughout the day from ~20 minutes before sunrise to ~20 minutes after sunset (https://www.timeanddate.com/sun/uk/maidstone) for three consecutive days. At these times the aid of artificial light was needed and so the first and last change over occurred in darkness. Fruit was offered throughout the night but was not changed. The period the fruit was deployed in the orchard was referred to as the ‘oviposition window’ as it was the period egg laying could have occurred. Data loggers (Lascar Electronics Ltd.) recorded temperature and humidity every 5 minutes throughout. Once removed from the station, samples were stored for 21 days at 23°C under a 12:12 light: dark (LD) cycle within a quarantine facility at NIAB EMR to allow eggs laid in the fruit to develop through to adult emergence. Specimen containers were sealed with a new ventilated lid to prevent asphyxiation of eggs within the fruit and subsequent offspring. Numbers of eggs were not counted as they could not be visually identified to species and, as wild unrestricted populations were used, other species might have deposited eggs on the fruit. Flies that emerged from the fruit were identified as either male *D. suzukii*, by the presence of spots on the wings and double sex combs on the front legs, female *D. suzukii*, by the oviscapts morphology, or other. *D. suzukii* were counted and here on are referred to as reproductive oviposition.

### Laboratory assays

*D. suzukii* cultures were established from a wild Italian strain collected in 2013. Populations of *D. suzukii* were housed in glass vials (Kimble Chase 25 × 95 mm Opticlear vials) at 23°C in a 12:12 L:D cycle with lights on at 08:00 and off at 20:00. Cultures and assays were housed within Percival DR-36VL environmental chambers within either the invertebrate facility at the University of Southampton or at NIAB EMR, Kent, UK. Chambers were programmed with a constant 65% humidity to prevent desiccation of media. Cultures were maintained on standard BDSC cornmeal food (100 % dH_2_O, 1 % Fisher agar, 9 % table sugar, 9 % precooked ground maize, 2 % baker’s yeast, 1 % soya flour, 5 % light spray malt, 0.3 % propionic acid, 0.3 % methyl benzoate dissolved in 3 % 70% ethanol (https://bdsc.indiana.edu/information/recipes/bloomfood.html) and were transferred into new vials every week. Cultures were kept genetically diverse by randomly mixing offspring to reduce genetic bottlenecks. As temperature and light cycles varied with experiment on commencing the assays, light and temperature cycles were modified within the chambers accordingly. For those conditions with a light phase over 12 hours a minimum of two chambers were used and were staggered to different ‘time zones’ enabling samples to be collected in a 12-hour time span and increasing replication. *D. suzukii* were acclimatised to experimental conditions for a minimum of 72 hours before starting assessments. Twenty-four hours before the start of laboratory trials, flies were removed from the chambers while in the photo phase. Three to seven-day old flies were immobilised on a CO2 pad (<u>Flystuff.com</u>) for a maximum of two minutes while sexed and transferred by the wing with soft forceps to experimental arenas, which varied with assay. In all assays’ it was presumed that flies had mated prior to the start of the experiment.

### Relationship between oviposition and adult emergence/ 12:12 LD 23°C

Groups of males and females were transferred to 70 ml specimen containers, as used in field oviposition assays, containing 10 ml of BDSC cornmeal media. Flies were subjected to constant 23°C under a 12:12 LD cycle. At lights on and lights off, flies were transferred to a new specimen container of media by tapping from the old container to the new. Eggs were counted under a microscope (x6 magnification) and the specimen container returned to the environment chamber. The same flies were transferred 6 times resulting in 3 days of assessments with 3 repetitions of light and dark counts. After 21 days the emerged adults were counted and egg to emergence survival assessed. The results from this assay were also used to deduce the light vs dark egg laying behaviour at a constant temperature (identified as 12:12 LD, 23°C in the results and discussion).

### Basic environmental cycle/17:7 LD 17/11°C

Groups of male and female *D. suzukii* were transferred to insect cages (17.5 × 17.5 × 17.5 cm, BugDorm-41515) housed within the environment chambers. Flies were subjected to a 17:7 LD cycle. Chambers were programmed with a day temperature of 17°C and a night temperature of 11°C with temperatures changing at lighting events. All cages were provisioned with water *ad libitum*, via a large Petri dish containing a soaked cotton wool reservoir, and food, a Petri dish (55 mm) of the BDSC cornmeal media. On commencing the trial, the Petri dish of media was replaced hourly for three days. Each dish was inspected under a microscope (x6 magnification) and the number of eggs counted.

### Recreated field conditions in the laboratory

Environment chambers were programmed with temperature cycles and lights-on and off times according to average field conditions during the reproductive oviposition field trials (June 11-22 °C, 18:6 L:D cycle 04:30 half-light, 05:00 full light, 21:30 half-light, 22:00 full-dark; August 14-32°C, 16:8 L:D cycle, 05:00 half-light, 05:30 full-light, 20:30 half-light, 21:00 full-dark; October 9-18 °C, 12.5:11.5 L:D cycle, 06:45 half-light, 07:15 full-light, 18:45 half-light, 19:15 full-dark). The maximum temperature used in the August conditions was reduced by 2°C due to avoid high heat-associated mortality observed in preliminary trials. At lights-on and off, half of the lighting banks in the chamber were changed 0.5 h prior to the other half to create a stepped transition. Groups of male and female *D. suzukii* were transferred to insect cages, as above, and housed within the environment chambers. All cages were provisioned with water *ad libitum* and a Petri dish (55 mm) of the BDSC media. On commencing the trial, the Petri dish of media was replaced every two hours for three days. Egg laying was assessed throughout the light phase only. Media was offered through the dark phase but was not changed or assessed. Each dish was inspected under a microscope (x6 magnification) and the number of eggs counted.

### Relationship between locomotion and oviposition in the laboratory

The Trikinetics Drosophila Activity Monitoring (DAM) system is commonly used to investigate locomotion rhythms in *Drosophila* ^21^. The DAM system sends infrared beams through the centre of a vial and monitors locomotion activity by analysing the number of infrared beam breaks in 5-minute periods or bins. Groups of 10 male and 10 female *D.suzukii* were loaded into LAM25 monitors (Trikinetics) in 25 mm × 95 mm glass vials containing 10 ml of a set sugar and agar food (dH_2_O, 5% sugar and 1% agar) sealed with a breathable cotton bung. Flies were taken from mixed sex cultures and were between 3- 7 days old at the start of the assay and presumed to have mated. Flies were immobilised and held on a CO2 pad before being transferred by the wing with soft forceps to the mouth of the vial. LAM25 monitors were housed in the Percival DR-36VL environment chambers under the conditions used in the three recreated field oviposition laboratory trial. Locomotion was monitored for a minimum of 6 days after a 24 hour ‘settling’ period.

### Statistical analysis

A linear regression analysis was performed to establish correlation between the number of eggs laid and subsequent adult emergence under laboratory conditions. Hourly oviposition in the laboratory and reproductive oviposition in the field was calculated to make results comparable across experiments. In those experiments when dark phases were assessed (12:12 LD 23°C, 17:7 LD 17/11°C in the laboratory and both field assays) Mann Whitney U test were performed for light vs. dark counts within each assay. Light counts/h were divided by dark counts/h to calculate the fold increase of light to dark oviposition rates. For both the August and October field and laboratory assays, normalised counts were calculated as a fraction of the total counts for each individual experimental day to allow for variation in population densities. A Mann Whitney U test was performed for pairwise comparisons of laboratory vs. field for normalised oviposition dependent on both time and temperature. Temperatures were divided into ranges and normalised oviposition analysed with a Kruskal-Wallis test to identify optimal temperature conditions. Statistical analysis was performed in SPSS.

For locomotion analysis, only groups in which all flies survived the full assessment period were used, resulting in a minimum of 6 groups per condition being analysed. Locomotion rates were analysed with ClockLab software through Matlab. Batch analysis was used to generate average locomotion activity from the sample groups. Average counts per minute and SEM were generated in ClockLab in 30-minute bins and combined into 2-hour intervals to match the format of the oviposition data.

## Results

### Correlation between oviposition and adult emergence

Due to technical constraints (see below) reproductive oviposition in the field was measured indirectly, as subsequent adult emergence. To help decide whether direct measurement of oviposition in the laboratory would be a suitable comparative measure of reproductive success the relationship of oviposition and subsequent *D. suzukii* adult emergence was examined under standard laboratory conditions (23°C 12h:12h LD). A linear relationship exhibited a high Pearson correlation coefficient (R^2^=0.938) and so adult emergence could be described as a linear function of oviposition and ~82% of eggs laid developed into adults (y= 0.8234) (Fig 1a). There was no significant difference in the correlation of egg to adult survival between oviposition that occurred during the light phase or in the dark phase. Thus, direct laboratory measurements of oviposition could be converted to estimated reproductive oviposition by applying a multiplication factor of 0.82.

**Fig 1a.**
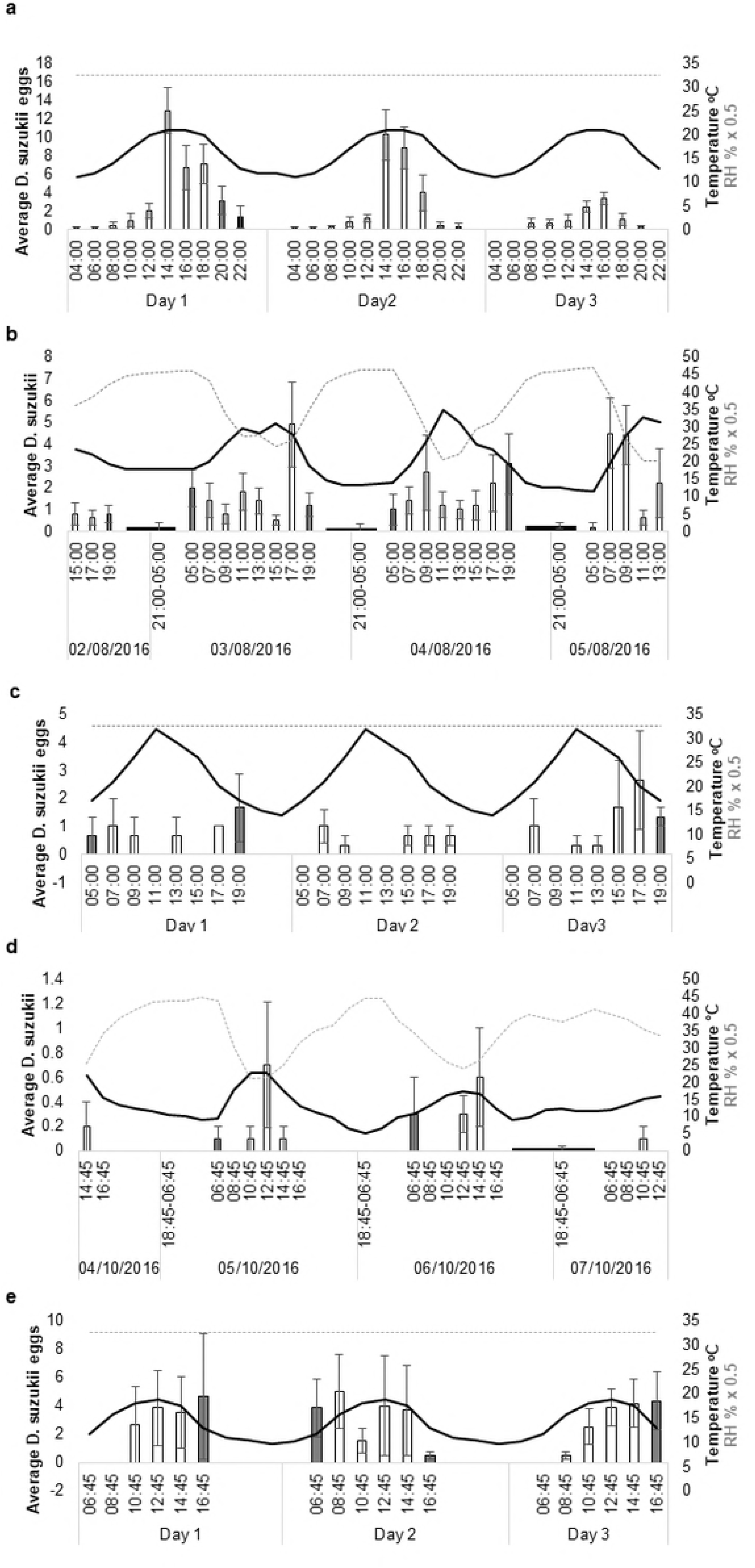
Correlation of number of eggs laid and adult emergence under 12:12 L:D cycle in a constant 23°C within the reproductive oviposition confirmation assay in the laboratory. Measurements during the day are indicated by an open circle, those laid during the night indicated by a closed triangle. 1b-e Average oviposition rate per hour (+SE) in either the day or night under (a)12:12 L:D in 23°C, (b) 17:7 17/11°C in the laboratory or reproductive oviposition rate per hour (+SE) in either the day or night under (c) August and (d) October in the field. * indicates significant difference between day and night counts within each environmental condtion

### Light versus dark

The average oviposition or reproductive oviposition per hour was significantly higher in the light compared to dark conditions in both laboratory and field assays. In 12:12 L:D cycle under constant 23°C 2.4-fold more eggs were laid in the light than in the dark (*p<*0.02) (Fig 1b). In the 17:7 LD 17/11°C conditions eggs laid per hour in the light were 15.4-fold higher than the night (*p*<0.000) (Fig 1c). In the August (Fig 1d) and October (Fig 1e) field experiments, hourly reproductive oviposition was 9.1- and 25-fold higher in the day than at night, respectively (both *p*<0.001).

### June conditions

The June field trial was excluded from the results and analyses due to insufficient field populations which resulted in no reproductive oviposition occurring. A late frost delayed blossom and cherry development which may have also affected population build up in the orchard at this time. Although fecund females were present in the days prior to the experiment it is likely that the population density was not sufficiently high enough to detect oviposition at this time. Environmental conditions were extrapolated from the field and re-created in the laboratory.

In the June laboratory assay there were consistent peaks in oviposition from 14:00 until 20:00 on each assessment day (Fig 2a). This coincided with peak temperatures between 19-22°C. In total, 571 eggs were laid across the three days. There were significant differences in normalised eggs in relation to both time and temperature (both *p*<0.000) (Fig 3a, b).

**Fig 2.**
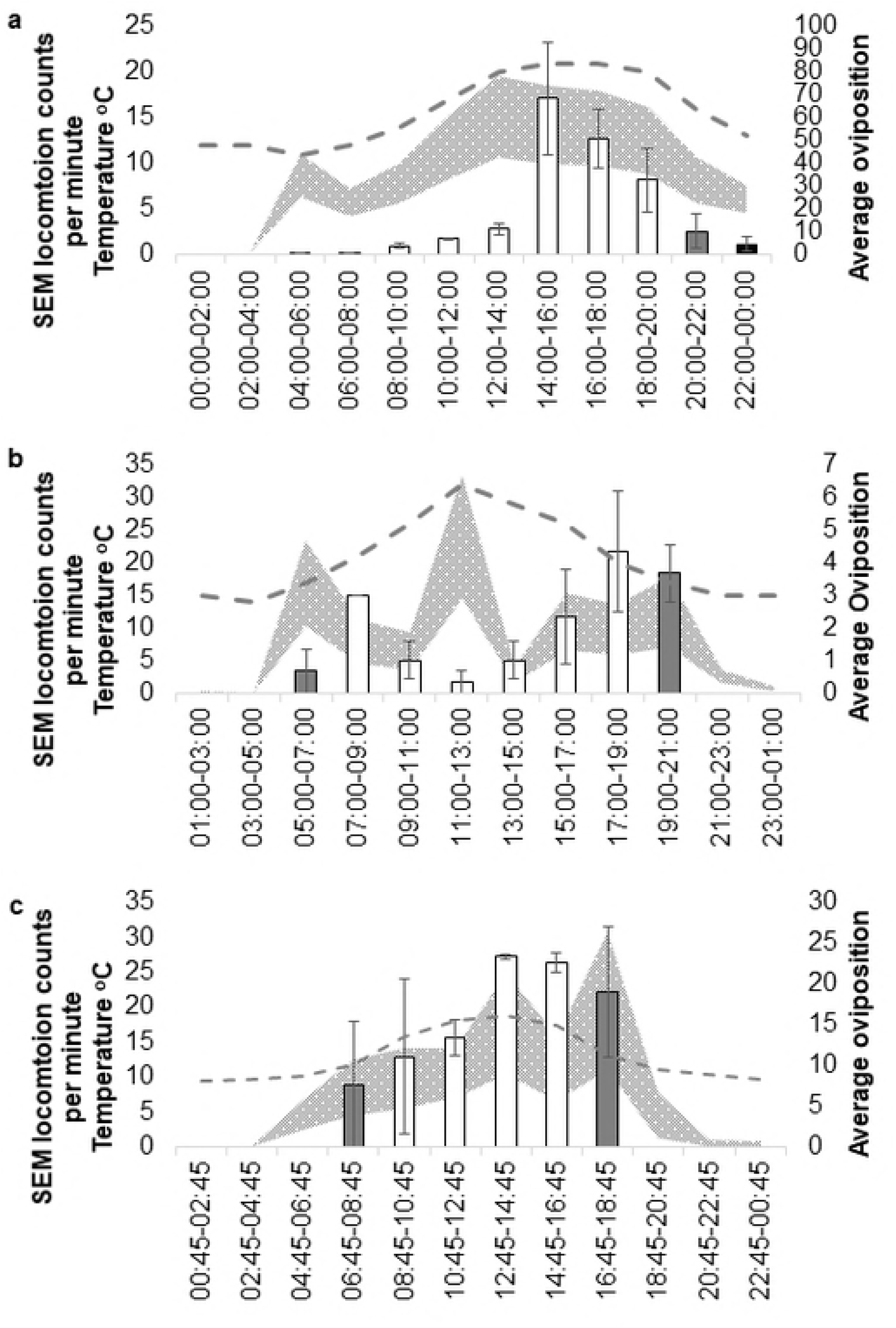
Average reproductive oviposition (+SE) from two hour ‘oviposition windows’ in August (b) and October (d) of wild *D. suzukii* in the field and average oviposition (+SE) (bars) and in recreated June (a), August (c) and October (e) conditions of laboratory strains in the laboratory. Average temperature (black solid line) and relative humidity (RH) (dashed grey line at x 0.5) over the three day. Note that relative humidity was kept constant at 65% in laboratory assays. Light is indicated by the colour of average *D. suzukii* bars. Black in darkness, grey in half light and white in light.

**Fig 3.**
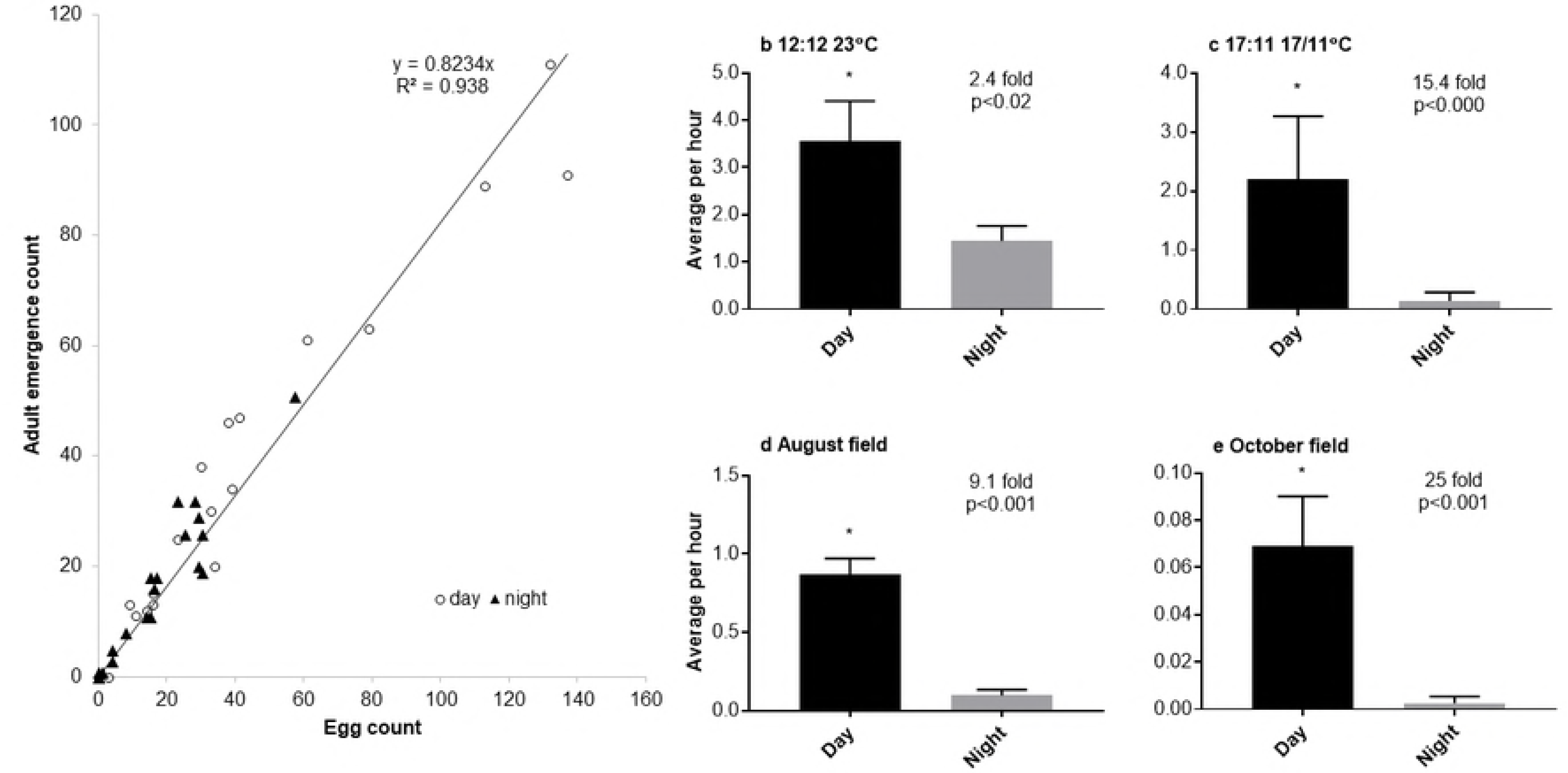
Normalised counts of *D. suzukii* laboratory oviposition (+SE) in June conditions in relation to time (a) and temperature (b).

### August conditions

During the August field assay, sunrise occurred at 05:24 and sunset at 20:42 on day one of the assay (2^nd^ August 2016). The photoperiod was 15.3:8.7 h light: dark. Reproductive oviposition occurred in two daily peaks on either side of the daily temperature warmest period (Fig 2b). The morning peak occurred 0-2 hours after sunrise on 3^rd^ August, 4-6 hours after sunrise on 4^th^ August and 2-4 hours after sunrise on 5^th^ August. Evening peaks occurred 12-14 hours after sunrise on 3rd August and 14-16 hours after sunrise on 4^th^ August with no obvious peak on 2nd August. Daytime dips in reproductive oviposition occurred when temperatures exceeded 30°C.

Although there was a strong diurnal rhythm in reproductive oviposition during the August field assay (see Fig 1d), no significant differences in normalised reproductive oviposition were found between the different time intervals during day light (Fig 4a). Nevertheless, there was a significant effect of temperature on normalised reproductive oviposition during day-light hours with highest normalised oviposition occurring between 27-30 °C (p<0.006) (Fig 4d).

**Fig 4.**
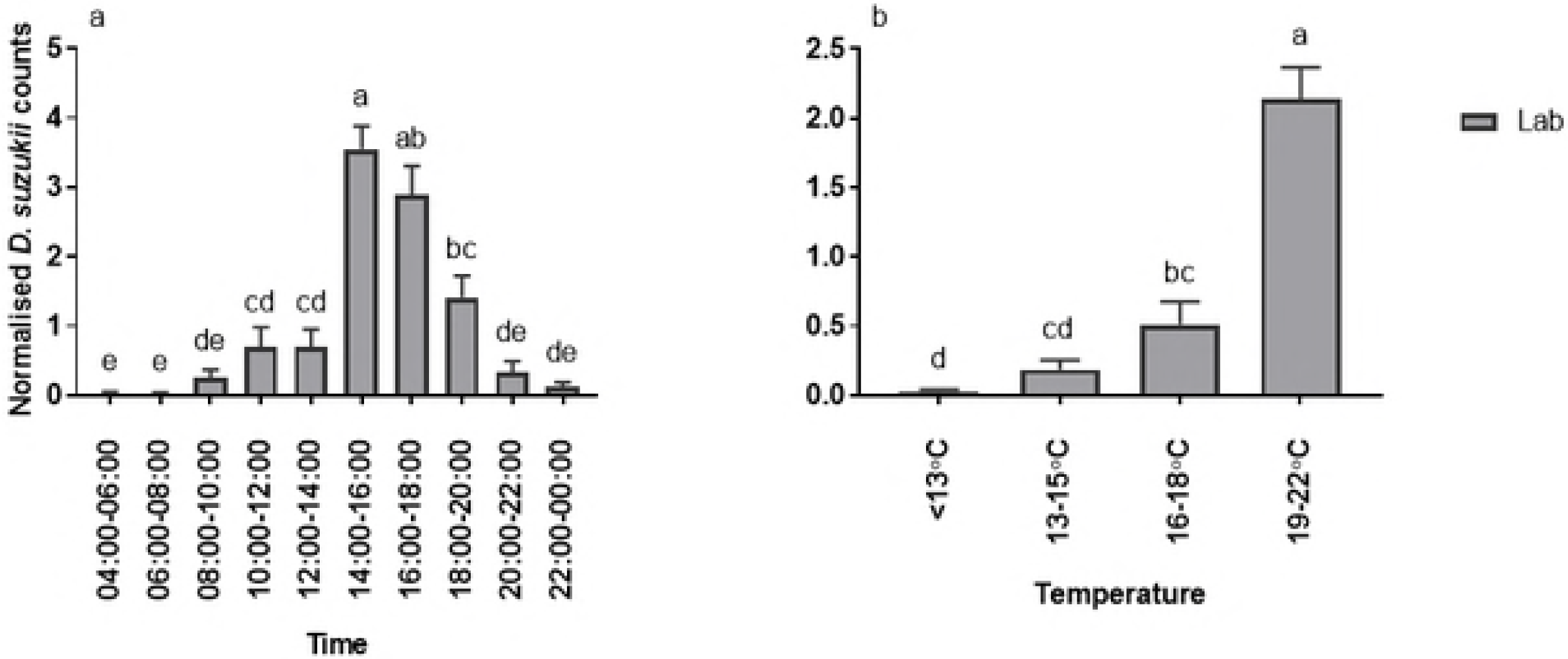
Normalised counts of *D. suzukii* reproductive oviposition from (a) field assay (+SE) (black bars) and (b) laboratory oviposition (+SE) (grey bars) in August conditions and (c) field vs. laboratory (+SE) in relation to time. Normalised counts of *D. suzukii* reproductive oviposition from (d) field assay (+SE) and (e) laboratory oviposition (+SE) in August conditions and (f) field vs. laboratory (+SE) in relation to temperature. Different letters identify significant differences between time points within each setting.

When August conditions were re-created in the laboratory, overall egg laying was low (49 in total) although egg laying did occur on each day of assessment (Fig 2c). There was a significant effect of time interval on normalised oviposition (*p*<0.011), (Fig 4b) with fewer eggs laid between 05:00-07:00, 09:00-11:00, 11:00-13:00 and 13:00-15:00. There was higher egg laying in temperatures between 19-22 °C although this was not significant (Fig 4e).

In the laboratory, peaks and troughs of *D. suzukii* oviposition, correlated with reproductive oviposition rates in the field. There was no significant difference between the laboratory and field counts in relation to time (Fig 4c). There was also no significant difference in counts between laboratory and field assays’ dependent on temperature ranges (Fig 4f). Regarding the apparent differences in the impact of daily time and temperature on day-time (reproductive) oviposition it should be noted that field temperature and humidity conditions were somewhat variable from day to day and that field temperatures were measured at the site of oviposition, which might have been visited only briefly by the flies.

### October conditions

In the October field assay, sunrise was at 07:04 and sunset at 18:28 on day one of the assay (5^th^ October 2016). The photoperiod was 11.4:12.6 L:D. Peaks in reproductive oviposition occurred 6-8 hours after sunrise on 5^th^ October and 8-10 hours after sunrise on 6^th^ October (Fig 2d). There was no obvious peak on the 7th October. Overall reproductive oviposition was low (25 in total) at this time in the field. Peaks in reproductive oviposition appeared to occur at peak temperature each day. There was a significant effect of time on normalised reproductive oviposition counts (*p*<0.039) (Fig 5a) with no reproductive oviposition occurring between 08:45-10:45 and 16:45-18:45 on any day. There was no significant effect of temperature on normalised reproductive oviposition (Fig 5d).

**Fig 5.**
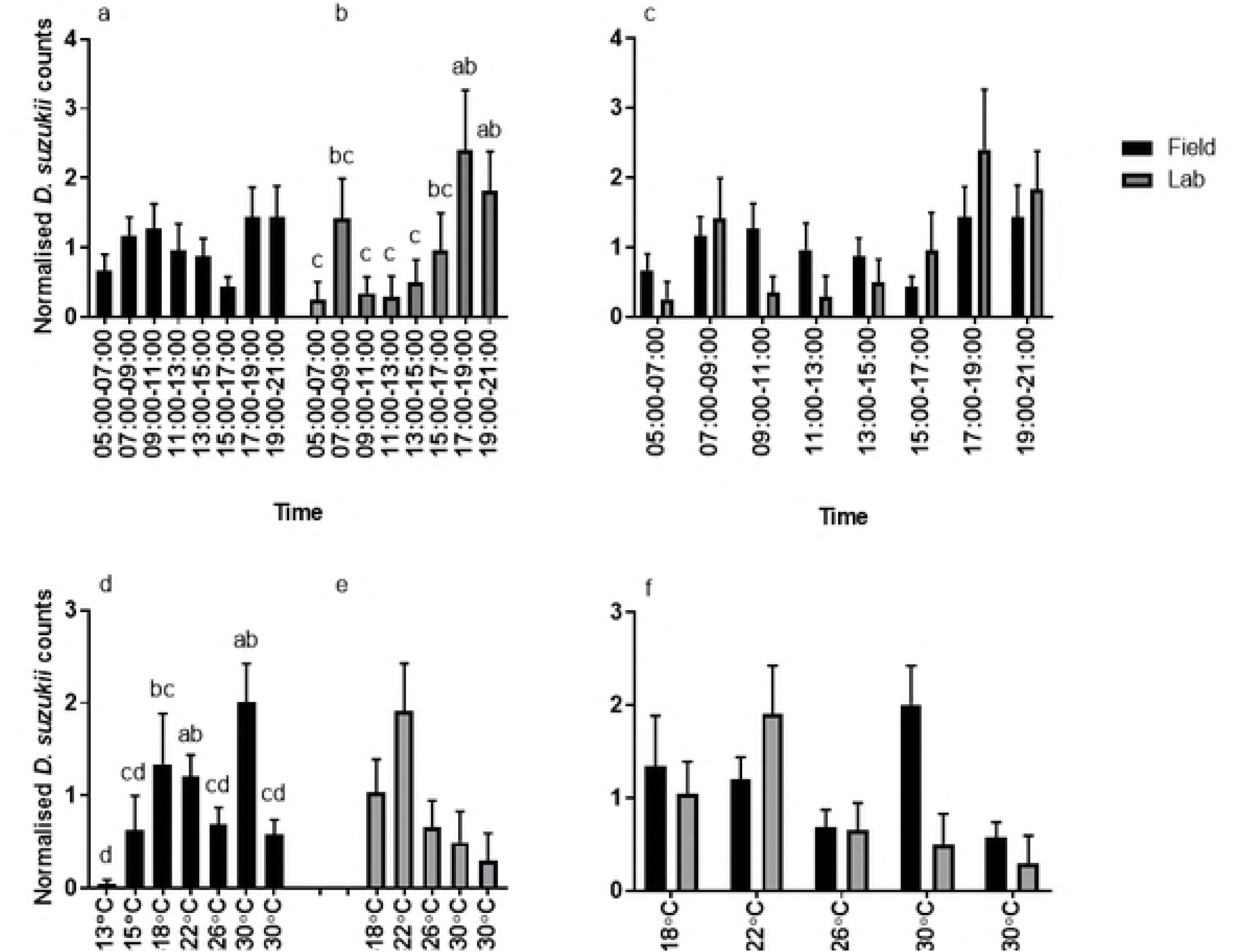
Normalised counts of *D. suzukii* reproductive oviposition from (a) field assay (+SE) (black bars) and (b) laboratory oviposition (+SE) (grey bars) in October conditions and (c) field vs. laboratory (+SE) in relation to time. Normalised counts of *D. suzukii* reproductive oviposition from (d) field assay (+SE) and (e) laboratory oviposition (+SE) in October conditions and (f) field vs. laboratory (+SE) in relation to temperature. Different letters identify significant differences between time points within each setting.

In the laboratory under the October conditions, oviposition occurred throughout each day with no specific peaks identifiable (Fig 2e). Overall 146 eggs were laid in total across the three days. There was no significant effect of time or temperature on normalised counts (Fig 5b and e).

There were significant differences between the laboratory and field assays with October conditions at only two time points; 08:45-10:45 (*p=*0.03) and 16:45-18:45 (*p=* 0.002). During these periods no eggs were laid in the field (Fig 5c). There were also significant differences in egg counts in 2 of the 4 temperature ranges (Fig 5f). At 13-15°C, counts in reproductive oviposition in the field were significantly lower than oviposition in the laboratory (*p*=0.05). However, at 16-18°C, counts in reproductive oviposition in the field were significantly higher than in the laboratory (*p*=0.02).

### Locomotion activity and oviposition

Locomotion activity profiles were collected under the same recreated temperature and light cycles used in the laboratory oviposition assay, obtained from the field conditions. Notably, times of peak locomotor activity coincided with peak oviposition for the June (Fig 6a) and October (Fig 6c), but not the August condition (Fig 6b). Further, oviposition rates at dawn in August and during the entire morning in June were lower than might be expected based on the accompanying locomotor activity levels. Generally, the best coherence between relative locomotor and oviposition activity levels was observed for the mid-afternoon to dusk period.

**Fig 6.**
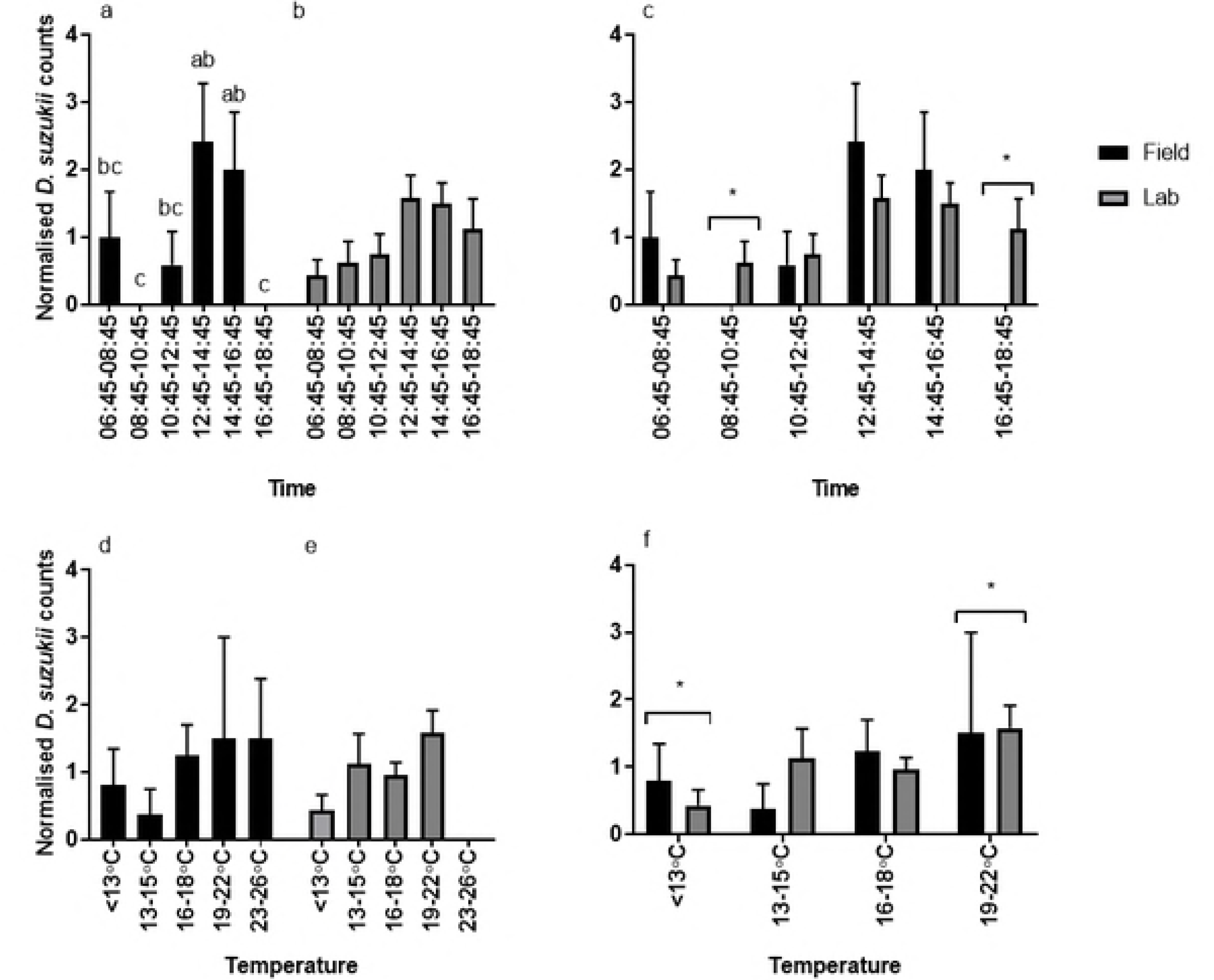
Mean ± SEM range of locomotion counts/minute of mix sex groups (grey area) in relation to average oviposition (+SE) (bars) and temperature (dash grey line) in the laboratory. Collected under recreated (a) June, (b) August and (c) October conditions in the laboratory. Light is indicated by the colour of average counts bars: Black in darkness, grey in half-light and white in full-light. Oviposition was assessed during the light phase (04:00-00:00 under June conditions, 05:00-21:00 under August and 06:45-18:45 under October).

## Discussion

In this study, diurnal reproductive oviposition rhythms were identified in wild *D. suzukii* populations that were then reproduced using simulated light and temperature profiles in the laboratory. As a result, we not only described reproductive oviposition rhythms of this horticultural pest in the field at relevant times of year, but also identified their environmental determinants. Furthermore, we show that laboratory-based experimental set-ups could be a valuable tool for predicting field behaviour across a wide range of environmental conditions.

We found a strong linear relationship between the numbers of *D. suzukii* eggs and the number of adult’s subsequently emerging following incubation under laboratory conditions. On average 82% of eggs survived to adult emergence justifying our approach of measuring daily patterns in reproductive success via reproductive oviposition in the field and via direct egg counts in the laboratory. Moreover, by normalising counts we were able to statistically test for differences in reproductive success profiles across the day in laboratory versus field conditions. Once collected from the field, all fruit and any eggs they contained were subjected to the same environmental conditions. Eggs laid during daylight hours in the field would have only been exposed to the variable environmental conditions in the orchard for the first 0- 2h interval. Therefore, we would expect mortality during development to be largely consistent throughout the different field samples.

In all trials, where reproductive success was compared between the light and the dark phase, significantly more (reproductive) oviposition occurred in the light. In a laboratory trial lacking temperature cues altogether (12:12 LD at constant 23°C) egg laying in the light was 2.5-fold higher than in the dark and this difference was strongly enhanced (15.4 fold) when a basic temperature cycle (17 °C during the light phase/11°C during the dark phase) was introduced along with a 17:7 LD cycle, all at constant relative humidity. Moreover, wild populations in the field exposed to natural fluctuations in light and temperature showed strong and significant day-time preferences for reproductive oviposition in August (9.1 fold) and October (25 fold).

In both August and October field trials, there was no consistent peak in reproductive oviposition in relation to sunrise, indicating that egg laying patterns are not solely reliant on lighting cues to entrain this behaviour. In the October field trial, peaks in reproductive oviposition occurred from oviposition windows that either encompassed peak temperature or 0-2 hours after. This was also found in the June and October laboratory trials.

In the August field trial, when fruit was deployed in oviposition windows in which temperature exceeded 30°C, reproductive oviposition was reduced within 0-2 hours. It is possible that there was a strict threshold for reproductive oviposition at temperatures above 30°C, but this could not be directly deduced from our data as temperatures were dynamic within the 2h interval surrounding temperature peaks.

Overall, lower reproductive oviposition occurred during the field trial in October versus the one in August, while the opposite was true for the corresponding laboratory assays. We attribute this apparent inconsistency to the relative population densities in wild habitats at these times of the year. *D. suzukii* not only exhibits reduced rates of growth and development when temperature decrease ^22^, it also transitions into a winter morph as females go into a reproductive diapause and do not oviposit through the colder, winter months ^23-25^. Winter morphs are produced as the result of larvae pupating in ~10°C and a short-day length ^26-28^. To examine whether female flies in the orchard had already entered diapause at the time of the October field trial, samples of flies trapped prior to the trial’s start were dissected. The presence of mature eggs across these samples indicated that oviposition in the October trial was not prevented by seasonal reproductive diapause.

Our observation of a strong day-time preference in reproductive oviposition occurred among both wild and laboratory populations and found the weakest preference for day-time oviposition under laboratory conditions at constant temperature. Our results are in apparent contrast to previous reports by Lin, et al. ^17^ and Evans, et al. ^18^ who both reported peaks in oviposition during the ‘dark’ phase. The former of these prior studies assessed oviposition under laboratory conditions (16:8 h LD 25°C) and found a significant increase for the ZT15.5-ZT19.5 interval (Zeitgeber Time with ZT0 corresponding to lights-on), whereas the latter examined oviposition in caged laboratory-raised *D. suzukii* under field conditions (~15:9 h LD with ~34°C day and ~24°C night) and found preferential oviposition for what amounts to the ZT14-ZT2. The discrepancies with our results may be explained by the warmer night temperatures in these studies and the inclusion of the period surrounding lights-off/dusk in their ‘night’ intervals. Our results for August field and laboratory conditions indicated that reproductive oviposition under these summer conditions is relatively high around dusk. Thus, separating dusk from night-time intervals may be critical in analyses that aim to predict the times of day when soft and stone fruit crops are at risk from *D. suzukii* oviposition particularly when extreme day temperature are reached.

Due to the modest counts of eggs or emerged adults per time point and the somewhat variable environmental conditions in the field, statistical analyses comparing different daytime intervals were limited in power. Nevertheless, we found that reproductive oviposition was significantly associated with time-of-day for the June and August laboratory and October field conditions and also with temperature for the August field and June laboratory conditions. Since egg laying is known to be under control of the circadian clock in *D.suzukii*’s sister species *D. melanogaster* ^29,30^, it is likely that the intrinsic daily timekeeping mechanisms in *D. suzukii* contributed to the observed changes in reproductive success between different times of day. The impact of environmental temperature on *D. suzukii* oviposition, has been directly demonstrated for laboratory-raised populations exposed to different constant temperatures ^22^ or in temperature gradients ^31^. Zerulla, et al. ^31^ investigated surface temperature of oviposition substrates and the effect on *D. suzukii* oviposition rates in the context of a temperature gradient and found a maximum oviposition rate at ~22°C. This is consistent with the optimal oviposition temperatures reported by prior studies comparing different constant temperature conditions ^22,32^ as well as the oviposition rates found for our laboratory trials mimicking June, August and October field conditions. Overall, 571 eggs were laid under the June conditions in the laboratory in comparison to the 49 and 146 laid for the August and October laboratory trials, respectively. Both October and August laboratory conditions included lower or upper development thresholds temperatures for *D. suzukii* ^33^ which is likely to have reduced oviposition rates within these experiments. Also, although flies were acclimatised to conditions for 72 hours before the laboratory trial commenced, it is possible that this was not long enough for laboratory strains to adapt to temperature ranges and could have promoted the low amount of oviposition ^34^. Within both the August laboratory and field trials, there was a reduction in reproductive success during time intervals with >30°C temperatures. Further, compared to the June and October conditions, peak reproductive oviposition was delayed from the daily temperature peaks in the early afternoon to relatively cooler temperatures around the onset of dusk. Moreover, day/night differences in field reproductive oviposition were lower during the August than the October trial. Taken together, these observations suggest that the daily profile of *D. suzukii* oviposition in the field is expected to reach its peak with the daily temperature maximum in the early afternoon. However, when temperatures exceed the preferred range for oviposition, the daily oviposition profile may become bimodal and exhibit daily maxima just before dusk. This hypothesis is further corroborated by the afore-mentioned results of Evans and co-workers ^18^, who found that in a day vs. night assay conducted in the context of high day-time temperatures (30-37°C), significantly more eggs were laid during the dusk-night-dawn interval than during core daylight hours. Temperatures of ≥30°C exceed the high temperature survival limit of *D. suzukii* ^22^ and may trigger escape behaviour to cooler locations if available. Indeed, Van Timmeren, et al. ^35^ observed a decrease in the number of wild *D. suzukii* on blueberry bushes during the warmest time of the day, when temperatures averaged ~30°C.

Our ability to reproduce daily patterns of reproductive oviposition in the field by recapitulating photoperiod and daily temperature gradient in the laboratory is illustrated for the August condition, where normalised reproductive oviposition counts exhibited no significant differences between laboratory and field assays. This was true both for pairwise comparisons per time interval and for temperature interval. Due to the sparse data under October field conditions no events were registered for some daily time intervals and these then emerged as times of day with significantly lower reproductive success rates in field versus laboratory. Significant differences between October laboratory and field conditions were also noted between normalised reproductive success at two intermediate temperature intervals. Nevertheless, the peaks in reproductive success during both field and laboratory October conditions coincided with the afternoon intervals and the temperature maxima. Thus, the laboratory trials for both August and October conditions successfully identified the daily peak in reproductive oviposition observed in the associated field trials. It is notable, that this was possible without reproducing the daily changes in humidity that occurred in the field. Humidity was kept constitutively high to avoid possible limitations of egg production and survival of both adults and offspring ^25,36^.

Since both oviposition ^17,18^ and locomotor activity ^11,37^ exhibit daily rhythmicity in *D. suzukii*, we explored the association between these two periodic behaviours under in identical environmental cycles in the laboratory. Although the relative rates of locomotor and oviposition activity seemed to be generally matched well in the mid-afternoon to dusk interval, locomotor activity spiked at other times without concomitant increases in egg laying. Increases in locomotor activity following lights-on for the June and August profiles resembled previous observations of immediate light-induced increases in locomotion under ‘summer’ conditions ^37^. It is not known to what extent these morning activity peaks depend on internal circadian clock function versus clock-independent light responses in locomotor activity, but the latter would not necessarily be functionally linked to oviposition. The locomotor activity peak observed around mid-day in the August experiment is reminiscent of high-temperature-associated afternoon peaks observed in *D. melanogaster* under semi-natural condition^12–16^ and may represent an escape response triggered by noxious heat. Consistent with this idea, this locomotor activity peak precisely coincides with the >30°C temperature peak. As *Drosophila* are ectotherms they do not maintain a constant body temperature and use the environment to regulate it, i.e. by moving into the sunlight when cold and moving to shade when hot ^38^. The increase seen in the mix sex group locomotion activity as temperatures reached 30°C could be the result of the flies increasing movement to try and locate a cooler niche. In the field, flies can escape from the rising temperatures and locate cooler areas to wait out the high heat. The resulting effect may be seen in both the laboratory and field oviposition trials as a reduction in oviposition as females are focusing on their own survival rather than egg laying. As eggs are immobile and unable to escape high temperatures, the reduction in reproductive oviposition in August could also be due to the females not depositing eggs in unfavourable environmental conditions and retaining eggs until conditions become optimum ^39^. Such retention of eggs by *D. suzukii* females until more favourable conditions occur has been termed ‘brood care’ ^31^.

## Conclusions

The main aims of this paper were to identify the daily oviposition rhythms of wild *D. suzukii*, promote these patterns in laboratory strains and identify vital environmental parameters required to do so. Although this is not the first report of patterns in *D. suzukii* oviposition behaviour, it is the first to document wild population oviposition behaviour under natural conditions in the UK in direct comparison with laboratory conditions. Our field results support theories suggested by other groups, that rhythmic patterns in oviposition are disrupted by unfavourable conditions; in this case extreme temperatures. With all the variations in factors between the laboratory and field assays including, substrate, humidity, lighting, setting and population strain, we were still able to promote field-like oviposition patterns in the laboratory. It therefore appears the correlation between the field and corresponding laboratory assay is dependent on temperature and photoperiod as the primary factors in influencing oviposition behaviour. From these results, it would be possible to make predictions of oviposition rhythms of wild *D. suzukii* populations from a laboratory-based experiment providing a realistic temperature cycle is used.

By comparing laboratory and field oviposition patterns with locomotion assays, we were also able to identify how these two behaviours interact with each other and temperature. The results of this paper highlight the importance of using realistic temperature and light cycles when investigating behavioural rhythm when field assays are not appropriate. By having a better understanding of how *D. suzukii* is behaving, dependent on light and temperature, means that potential control measures can be tailored to exploit these patterns.

## Acknowledgments

Thank you to G.H. Dean, Norton Folgate and Driscoll’s for fruit donations for this research. Thanks to the NIAB EMR, University of Southampton staff and the UK Spotted Wing Drosophila Working Group for support with this work.

